# EPHA4 Regulates the Transition between Mesenchymal and Hybrid States in TNBC

**DOI:** 10.64898/2026.10.04.756472

**Authors:** Wenchao Liu, Hoi Wan Lee, Madeline H. Bungert, Long C. Nguyen, Aleksandra Kurowska, Thomas Jiyoung Li, Geetha P. Yerradoddi, Eva Suarez, Marsha Rich Rosner

**Author notes:** Correspondence: Marsha Rich Rosner. These authors contributed equally to the work (Wenchao Liu, Hoi Wan Lee).

## Abstract

Metastatic progression in triple-negative breast cancer (TNBC) is strongly influenced by epithelial-mesenchymal plasticity (EMP), which enables transitions between epithelial, hybrid epithelial/mesenchymal (E/M), and mesenchymal cell states. While mesenchymal cells are highly invasive and resistant to therapy, hybrid E/M states confer advantages during metastatic colonization. Both mesenchymal and hybrid states therefore contribute to tumor aggressiveness through distinct mechanisms. However, the molecular regulation governing transitions between mesenchymal and hybrid E/M states remains poorly understood. Here, we identify EPHA4 as a regulator of this transition in TNBC. EPHA4 expression is enriched in mesenchymal TNBC cells, and genetic knockout of EPHA4 in MDA-MB-231 xenograft and 4T1 syngeneic models induces a shift toward hybrid E/M phenotypes. This transition reduces cellular invasion while enhancing metastatic colonization, consistent with the clinical observations of decreased EPHA4 expression in secondary metastases compared to primary tumors. Despite compensatory changes in other Eph/ephrin family members following EPHA4 loss, EPHA4 deficiency is sufficient to drive this phenotypic shift. These findings identify EPHA4 as a regulator of cancer cell plasticity in TNBC and highlight the therapeutic complexity of targeting pathways that govern metastatic state transitions.

## Introduction

Metastasis accounts for the majority of breast cancer-related deaths, yet limited insight into its regulation has impeded development of anti-metastatic treatments. Triple-negative breast cancer (TNBC) is the most aggressive breast cancer subtype but lacks effective targeted therapies. One promising direction for new treatments relates to cell state transitions. The epithelial-mesenchymal transition (EMT) is a critical developmental process that, when dysregulated, drives cancer progression (1, 2). Most research has focused on the initial transition from epithelial cells to mesenchymal states, or vice versa. While EMT was previously viewed as a binary switch, it is now recognized that cancer cells often occupy intermediate “hybrid E/M” states, expressing both epithelial and mesenchymal markers, a phenomenon known as epithelial-mesenchymal plasticity (EMP) (1–3). Mesenchymal cells are highly invasive and therapy-resistant, while hybrid E/M cells have higher metastatic potential (1, 2, 4). Complete reversal of EMT has been explored as a potential approach for reprogramming cancer cells back to a benign state, yet partial reversal to a hybrid state could lead to undesired effects (5). The mechanism by which fully mesenchymal cells can be reprogrammed to hybrid E/M states is not well understood.

Reprogramming of cell-cell interactions plays an important role in EMP cell state transitions (4, 6). Among the proteins that control these interactions in development are members of the Eph/Ephrin family (7). In this study, we investigated whether EPHA4 might contribute to epithelial-mesenchymal plasticity in breast cancer. Using patient tumor datasets, transcriptomic profiling, and functional studies across xenograft and syngeneic TNBC models, we show that EPHA4 is highly expressed in mesenchymal cells, particularly in TNBC (8, 9).

Genetic knockout of EPHA4 promotes a shift from mesenchymal toward hybrid E/M states, reducing invasive capacity while enhancing metastatic colonization. These findings identify EPHA4 as a regulator of cancer cell EMP and highlight the context-dependent complexity of targeting pathways that govern metastatic state transitions.

## Results

### Section 1. EPHA4 is highly expressed in mesenchymal-like breast tumors

To identify Eph/Ephrin genes that are preferentially expressed in mesenchymal versus hybrid EMP cell states in breast cancer, we analyzed transcriptomic data from two treatment-naïve patient cohorts: TCGA-BRCA, representing a diverse patient population, and ISPY2, consisting of higher-risk, aggressive breast cancer cases (10) (Fig. 1a). Using established epithelial and mesenchymal marker genes (11), we classified tumors as epithelial, hybrid, or mesenchymal. Notably, the ISPY2 cohort contained a higher proportion of E/M hybrid tumors compared to TCGA, reflecting its enrichment for high-risk patients (Fig. 1b).

**Figure 1.**
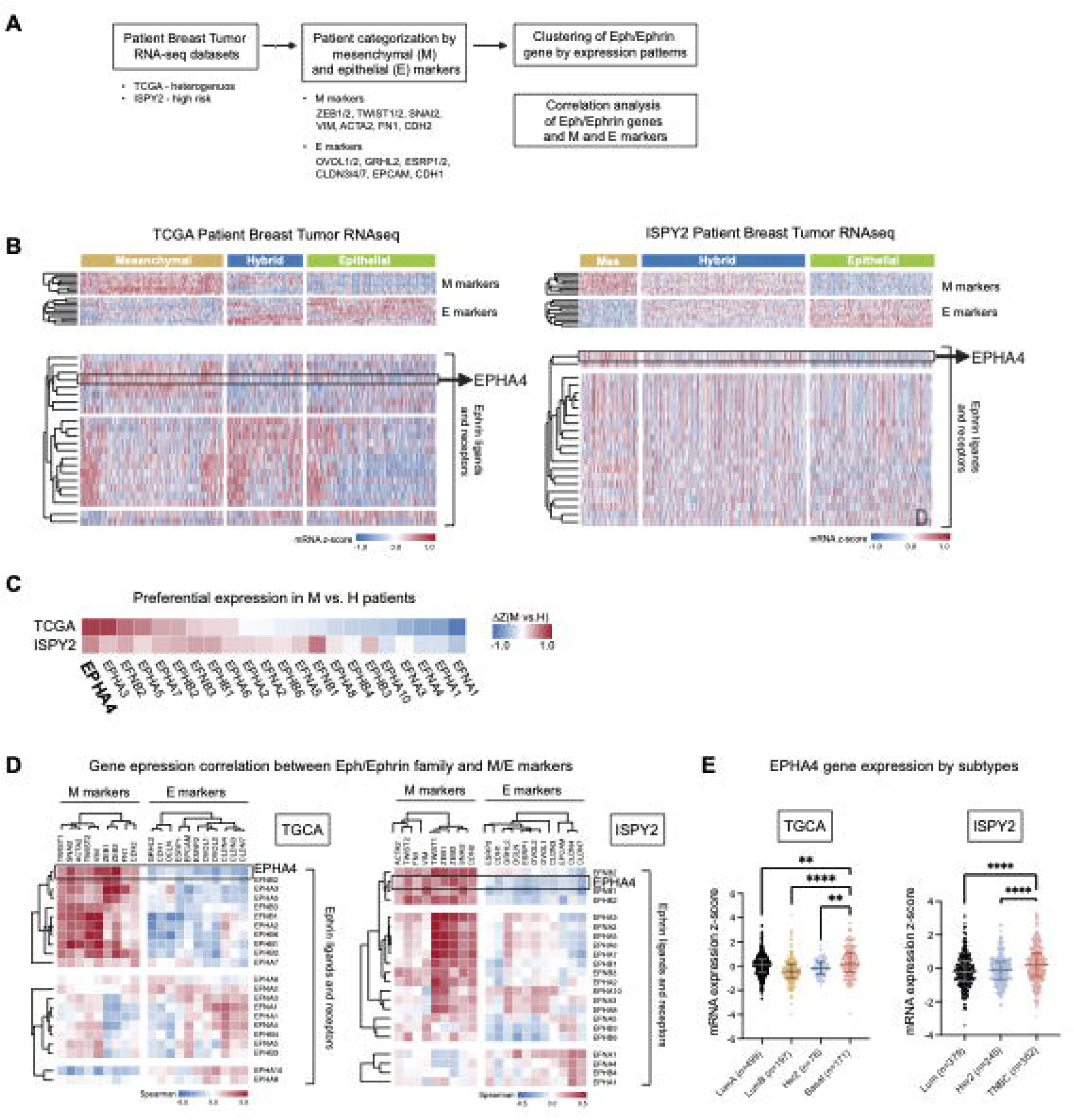
EPHA4 is enriched in mesenchymal-like breast tumors. **(A)** Patient data analysis workflow. RNA-seq data from TCGA-BRCA and ISPY2 breast tumors were stratified into epithelial (E), hybrid (H), and mesenchymal (M) states based on established marker genes, followed by analysis of Eph/ephrin gene expression and correlation with E and M markers. **(B)** Heatmaps showing expression of epithelial and mesenchymal marker genes (top) and Eph/ephrin ligands and receptors (bottom) across tumors grouped by epithelial-mesenchymal state. Color scale represents mRNA z-score. EPHA4 is highlighted. Left, TCGA dataset; right, ISPY2 dataset (GSE194040). **(C)** Preferential expression of Eph/ephrin genes in mesenchymal versus hybrid tumors in TCGA and ISPY2 datasets. Color scale represents ΔZ (M vs. H) = z-score(M) - z-score(H). Genes ranked by ΔZ (M vs. H) in TCGA. **(D)** Correlation between Eph/ephrin genes and epithelial or mesenchymal marker genes in TCGA (left) and ISPY2 (right). Color scale indicates spearman correlation coefficient. **(E)** Relative expression of EPHA4 in breast tumors of various molecular subtypes in TCGA (left) and ISPY2 (right).

Across the Eph/Ephrin family, EPHA4 showed the most consistent preferential expression in mesenchymal tumors in both datasets, with strong positive correlations to mesenchymal markers and negative correlations to epithelial markers (Fig. 1b-d). Further analysis by molecular subtype demonstrated that basal-like primary tumors, the most aggressive subgroup, displayed the highest levels of EPHA4 expression (Fig. 1e). Expression analysis of breast cancer cell lines (12) further confirmed that EPHA4 is enriched in mesenchymal TNBC cells (Fig. S1). Based on these findings, we initially selected the human MDA-MB-231 cell line, which exhibits a mesenchymal phenotype and high endogenous EPHA4 levels, to investigate EPHA4’s functional role in this aggressive tumor subset.

### Section 2. Loss of EPHA4 induces mesenchymal to E/M hybrid TNBC cell state transitions

To investigate EPHA4’s role in mesenchymal TNBC tumors, we used CRISPR-Cas9 to knock out EPHA4 in MDA-MB-231 cells and performed bulk RNA-seq on xenograft tumors (Fig. 2a; Fig. S2a). Differential gene expression and functional enrichment analysis identified “cell junction organization”, a key epithelial process, as the most significantly upregulated pathway in EPHA4 knockout (KO) tumors (Fig. 2c; Fig. S2b, c). We observed a marked increase in epithelial markers and cell-cell junction components (Fig. 2b), which was confirmed by qPCR (Fig. 2b). In contrast, mesenchymal markers showed variable regulation, with most being downregulated (Fig. S2d, e). Overall, these findings demonstrate that EPHA4 loss in mesenchymal TNBC cells promotes a hybrid epithelial/mesenchymal (E/M) gene expression state.

**Figure 2.**
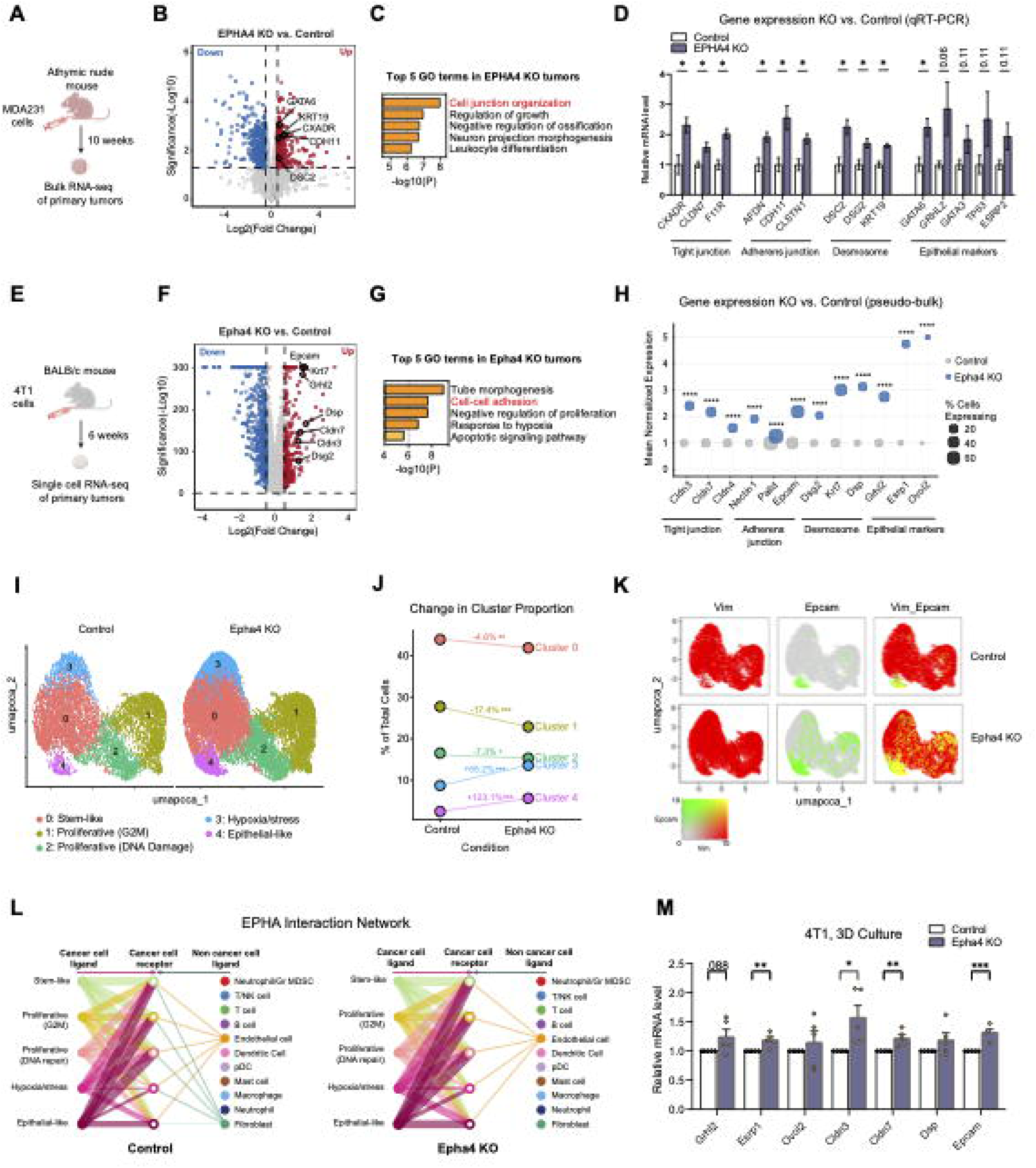
Loss of EPHA4 promotes epithelial gene expression and hybrid cell states in TNBC models. **(A)** Experimental design of MDA-MB-231 xenograft tumor bulk RNA-seq experiment. **(B)** Differentially expressed genes comparing EPHA4 KO to control MDA-MB-231 tumors (n = 4 tumors for control; n = 5 tumors for EPHA4 KO). Genes involved in cell-cell adhesion are highlighted. **(C)** Metascape enrichment analysis of genes significantly upregulated (Log2FC > 0.5, p < 0.05) in EPHA4 KO versus control MDA-MB-231 tumors. **(D)** qPCR validation of cell-cell junction and epithelial marker gene expression in EPHA4 KO versus control MDA-MB-231 tumors (n = 4 tumors for control; n = 5 tumors for EPHA4 KO). Data represent mean ± SEM. Two-tailed Mann-Whitney test, p < 0.05*. **(E)** Experimental design of 4T1 syngeneic tumor scRNA-seq experiment. **(F)** Pseudo-bulk analysis of differentially expressed genes in Epha4 KO versus control 4T1 tumor cells (n = 2 tumors for control; n = 2 tumors for Epha4 KO). Cell-cell adhesion genes and epithelial marker genes are highlighted. **(G)** Metascape enrichment analysis of genes significantly upregulated (Log2FC > 0.5, adjusted p-value < 0.05) in Epha4 KO versus control 4T1 tumor cells. **(H)** Pseudo-bulk expression of cell-cell junction and epithelial marker genes in control and Epha4 KO tumor cells. Dot size indicates fraction of expressing cells; y-axis shows mean normalized expression. **(I)** UMAP visualization of 4T1 tumor cells from control and Epha4 KO tumors, colored by cell state clusters. **(J)** Changes in cluster proportions between control and Epha44 KO tumors. **(K)** Expression of mesenchymal marker (Vim) and epithelial marker (Epcam) across tumor cell clusters in control and EPHA4 KO tumors. Co-expression is shown in yellow. **(L)** Inferred EPHA signaling network within the tumor microenvironment using CellChat. Interactions of cancer cell receptors with ligands from cancer cell subclusters or stromal/immune populations are shown. **(M)** qRT-PCR analysis of epithelial gene expression in control and EPHA4 KO 4T1 cells cultured in 3D Matrigel for three days. Data represent mean ± SEM from five independent experiments. Two-tailed student t-test, p < 0.05*, p < 0.01**, p < 0.001***.

To assess EPHA4’s role in cancer cells within an immune-competent tumor microenvironment, we generated Epha4 knockout tumors using the 4T1 mesenchymal TNBC syngeneic mouse model (Fig. 2e; Fig. S3a). Single cell RNA-seq revealed a heterogeneous tumor microenvironment (Fig. S3b). Similar to the MDA-MB-231 results, pseudobulk analysis of 4T1 cancer cells showed that EPHA4 deficiency significantly upregulated cell-cell adhesion pathways and epithelial genes (Fig. 2f-h; Fig. S3c, d-left), while mesenchymal gene levels were largely unchanged (Fig. S3d).

Further subclustering identified five cancer cell phenotypes: proliferative [G2M], proliferative [DNA repair], hypoxia/stress, stem-like, and epithelial-like (Fig. 2i; Fig. S3w). We observed a marked increase in the proportion of cells in the epithelial-like and hypoxic stress clusters (Fig. 2j). Importantly, while epithelial gene expression was limited to the epithelial-like cluster in control tumors, Epha4 KO tumors exhibited widespread acquisition of epithelial markers across additional clusters, including those with strong mesenchymal features, indicative of a shift towards hybrid epithelial/mesenchymal cell states (Fig. 2k; Fig. S3f).

EPHA4-mediated juxtracrine signaling is activated through interactions with membrane-bound ephrin ligands (types A and B) (13). To elucidate the cell-cell interactions regulating EPHA4 function within tumors, we analyzed EPHA and EPHB signaling networks by CellChat (14). Both pathways were primarily active in tumor cells, with minor contributions from endothelial and fibroblast populations (Fig. 2l; Fig. S3g). Contrary to previous reports (15), we did not detect significant ligand input from macrophages or other immune cells in this model. Supporting these *in vivo* findings, EPHA4 knockout in 4T1 cells grown in a 3D Matrigel environment, but not in 2D culture, resulted in upregulation of epithelial marker genes compared to controls (Fig. 2m; Fig. S3h). These data suggest that EPHA4 predominantly represses epithelial gene expression through tumor cell-intrinsic mechanisms, which are dependent on a 3D microenvironment.

### Section 3. Loss of EPHA4 reduces invasion yet enhances lung colonization burden in TNBC

Having observed the change in EMP gene expression, we looked for phenotypic changes corresponding to a shift from mesenchymal to hybrid cell states. Compared to the mesenchymal state, hybrid E/M states have been associated with reduced invasiveness but increased metastatic potential (4). In MDA-MB-231 cells, EPHA4 knockout did not affect primary growth in athymic nude mice (Fig. 3a, b). Due to inefficient spontaneous metastasis (16), we assessed invasion and metastatic colonization directly. EPHA4 knockout reduced invasion *in vitro* (Fig. 3c), consistent with prior siRNA studies (17), but significantly increased lung metastatic colonization, as indicated by larger colonies (Fig. 3d-f).

**Figure 3.**
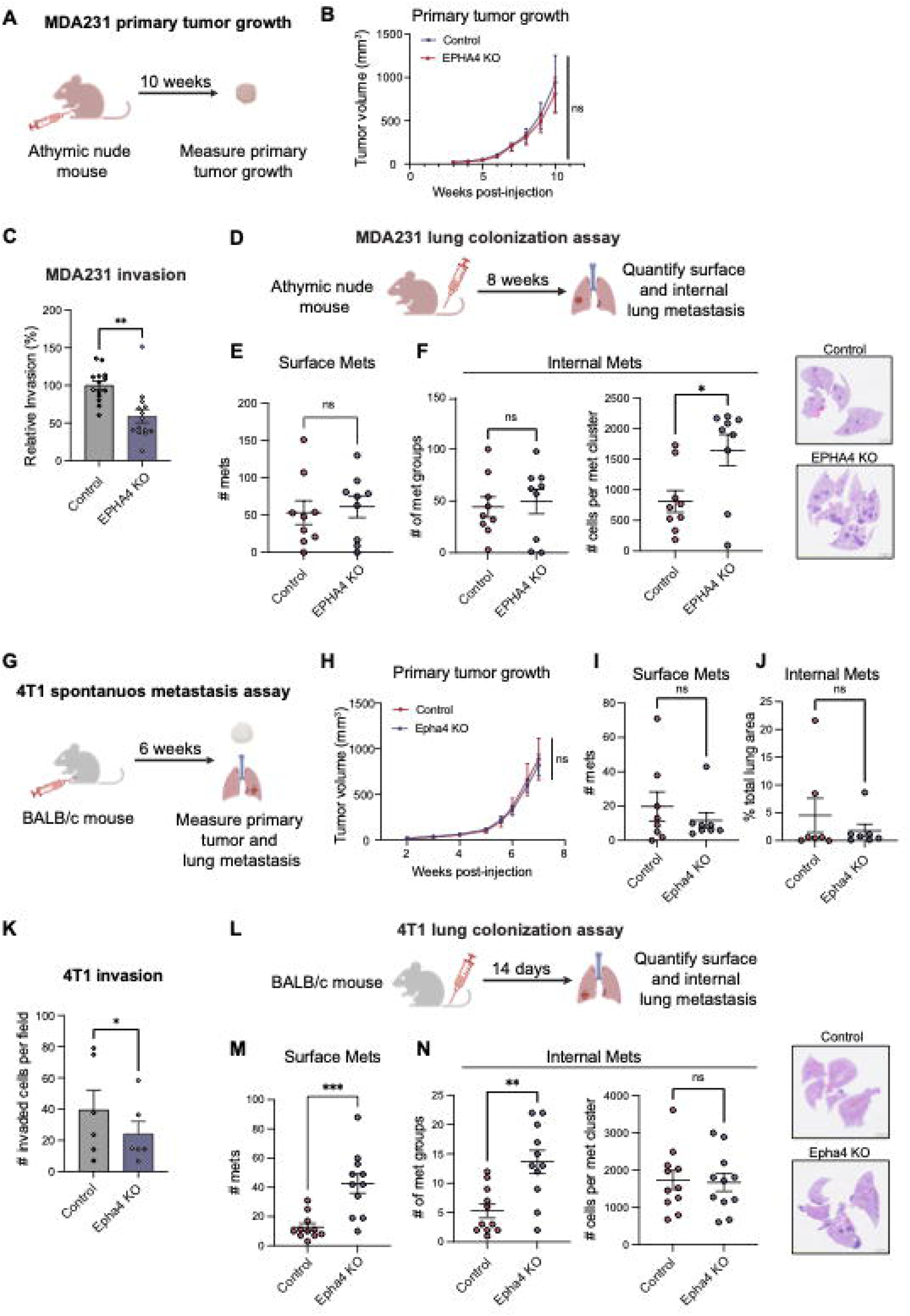
Loss of EPHA4 reduces invasion but enhances lung colonization in TNBC models. **(A)** Experimental design for MDA-MB-231 xenograft primary tumor growth in athymic nude mice. **(B)** Primary tumor growth of control and EPHA4 KO MDA-MB-231 tumors. Tumor volume was measured by caliper. **(C)** *In vitro* transwell invasion assay of MDA-MB-231 control and EPHA4 KO cells. Data represent mean ± SEM from four independent experiments. Two tailed, ratio-paired t test, p < 0.01**. **(D)** Experimental design of MDA-MB-231 lung colonization assay in athymic nude mice. **(E)** Surface macro-metastasis counts from MDA-MB-231 lung colonization assay. N = 9 mice for control; n = 9 mice for EPHA4 KO. **(F)** Quantification of internal lung metastases, including number of metastatic foci and cells per metastatic cluster. Representative H&E-stained lung sections are shown. **(G)** Experimental design of the 4T1 spontaneous metastasis assay in BALB/c mice. **(H)** Primary tumor growth of control and Epha4 KO 4T1 syngeneic tumors. Tumor volume was measured by caliper. N = 8 tumors for control; n = 8 tumors for Epha4 KO. **(G)** Surface macro-metastasis counts from 4T1 spontaneous metastasis assay. N = 8 lungs for control; n = 8 lungs for Epha4 KO. **(H)** Quantification of internal metastasis burden by H&E staining (one slide section per mouse). N = 7 lungs for control; n = 7 lungs for Epha4 KO. **(I)** *In vitro* transwell invasion assay of 4T1 control and Epha4 KO cells. Results from three independent experiments. **(J)** Experimental design of the 4T1 control and Epha4 KO cell lung colonization assay in BALB/c mouse. **(K)** Surface macro-metastasis counts from 4T1 lung colonization assay. N = 9 mice for control; n = 9 mice for Epha4 KO. **(L)** Quantification of internal lung metastases, including number of metastatic foci and cells per metastatic cluster. Representative H&E-stained lung sections are shown. For statistical test in all panels: two-tailed student t-test, p < 0.05*, p < 0.01**, p < 0.001***.

In the 4T1 model, EPHA4 knockout did not significantly alter primary tumor growth or total spontaneous metastasis (Fig. 3g-j). However, when assessed directly, EPHA4 loss similarly decreased invasion (Fig. 3k) and potentiated lung colonization through increased lung colony number (Fig. 3l-n); these opposing effects likely explain the lack of apparent change in spontaneous metastasis. Overall, loss of EPHA4 shifts cancer cells from a highly invasive mesenchymal state to an E/M hybrid state with enhanced metastatic colonization.

### Section 4. EPHA4 is a cancer-intrinsic driver of the hybrid to mesenchymal cell state transition

Ephrin ligands and receptors form a redundant family, enabling compensation when one member is lost (18–21). Following EPHA4 knockout, we observed distinct upregulation of other ephrin genes: EPHA2, EPHB1, and EPHB6 in MDA-MB-231 tumors, and EFNB1, EFNA1, EPHB2, and EPHB6 in 4T1 tumors (Fig. 4a). qPCR confirmed compensatory gene induction in MDA-MB-231 tumors (Fig. S4a). *In vitro*, compensatory genes were not upregulated in either cell type (Fig. S4b, d), suggesting that compensation is driven by the tumor microenvironment rather than by intrinsic cancer cell pathways. In patient tumors, compensatory gene expression did not consistently associate with hybrid EMP states (Fig. 1c). Although post-transcriptional compensation cannot be excluded, these results suggest that EPHA4 is the primary driver of cancer cell-intrinsic EMP reprogramming rather than acting via compensatory pathways.

**Figure 4.**
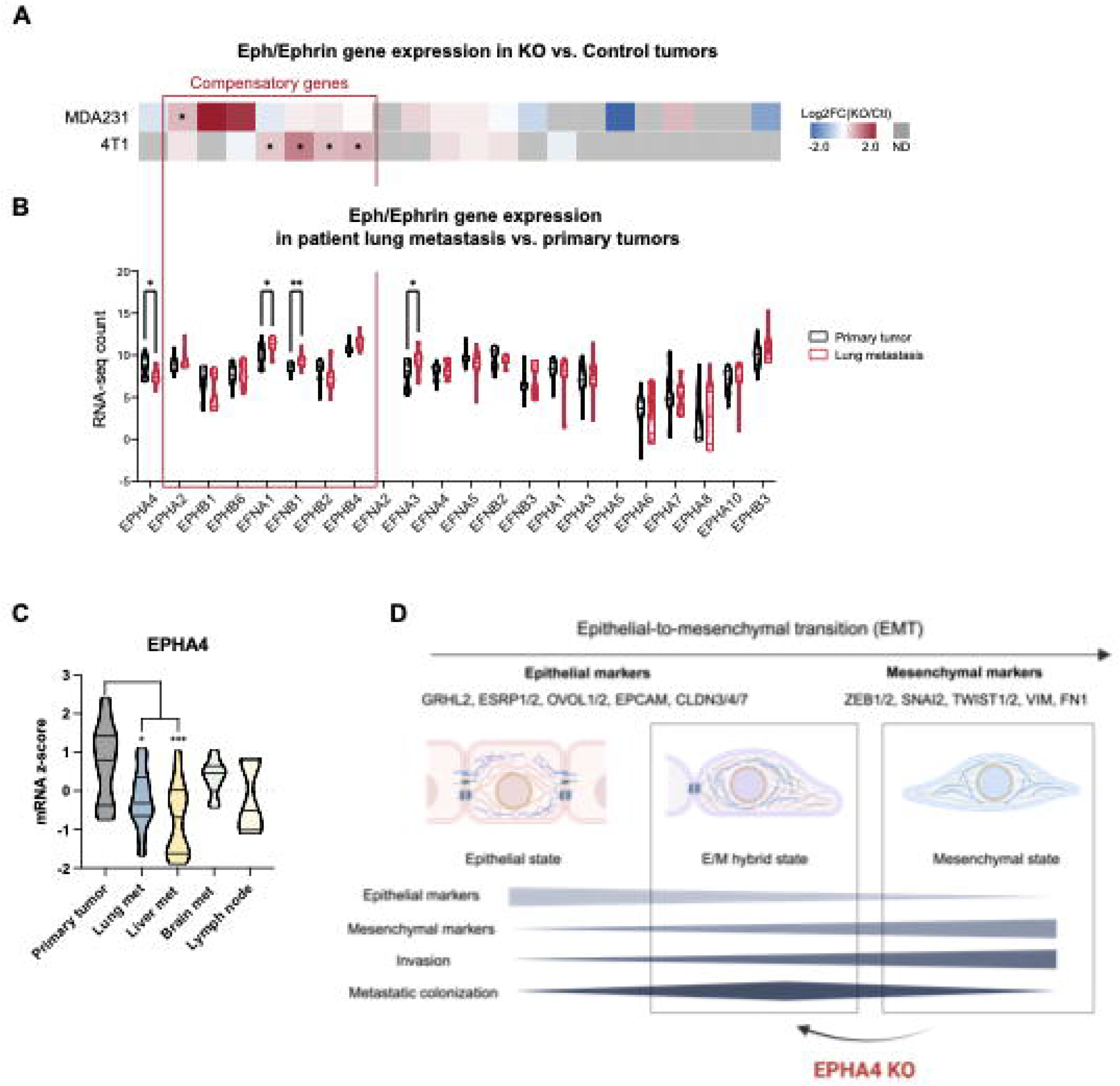
Loss of EPHA4 led to compensation within the Eph/Ephrin gene family. **(A)** Heatmap showing changes in Eph/ephrin gene expression in EPHA4 KO versus control tumors in MDA-MB-231 and 4T1 models. Genes that were significantly upregulated in EPHA4 KO tumors are marked with stars. Other near significant p-values are indicated. Color scale represents log2 fold change (KO/control). **(B)** Expression of Eph/ephrin genes in matched primary breast tumors and lung metastases from patients (16 patients, GSE110590). **(C)** Relative EPHA4 expression (mRNA z-score) across matched primary breast tumors and metastatic sites. Data also from GSE110590. **(D)** Schematic model illustrating the role of EPHA4 in regulating epithelial-mesenchymal plasticity (EMP). Loss of EPHA4 shifts cells toward a hybrid E/M state associated with reduced invasion and enhanced metastatic colonization. Illustration adapted from review (4).

Given that EPHA4 promotes invasion but restrains metastatic colonization, we examined its expression in matched patient samples. EPHA4 was the only ephrin receptor gene reduced in lung metastases (Fig. 4b) and was consistently downregulated in secondary metastatic sites compared to primary breast tumors (Fig. 4c). By contrast, EFNA1 and EFNB1, two compensatory genes upregulated in EPHA4 KO 4T1 tumors that have been associated with metastatic progression (22, 23), were elevated in breast lung metastases (Fig. 4b), potentially contributing to the enhanced colonization seen in the 4T1 model.

## Discussion

In this study, we show that EPHA4 is a regulator of mesenchymal cell state stability in aggressive breast cancer. Loss of EPHA4 in mesenchymal TNBC cells induces a transition toward hybrid epithelial/mesenchymal (E/M) states, accompanied by reduced invasive capacity but enhanced metastatic colonization. These findings align with prior studies identifying EPHA4 as a downstream target of established EMT regulators, including Twist1(24) and TGFβ(9), in development and breast cancer. Given the association between mesenchymal states and drug resistance(4), our results also have therapeutic implications. Although EPHA4 and related signaling pathways may represent attractive targets in aggressive cancers due to their association with mesenchymal phenotypes and invasion, our data suggest that inhibition of EPHA4 alone may produce unintended consequences by promoting transitions toward more metastatic, hybrid E/M states. This interpretation is supported by clinical observations showing reduced EPHA4 expression in metastatic lesions relative to primary tumors.

EPHA4 is best known as a receptor tyrosine kinase that mediates cell-cell repulsion through post-translational regulation of adhesion proteins (25). Here, we found that EPHA4 also suppresses cell-cell adhesion molecules at the transcriptional level. Our results raise the possibility that the balance between cell-cell repulsion and adhesion, regulated by ephrin ligands and receptors, could play a critical yet underappreciated role in cancer progression by regulating transitions between diverse EMP cell states.

Our study found an apparent functional tradeoff between invasion and metastatic colonization following EPHA4 loss, consistent with the emerging concept that the hybrid E/M states confer greater metastatic fitness than the mesenchymal state that drives invasion. EPHA4 loss in 4T1 tumors, when compared to MDA-MB-231 tumors, induced stronger epithelial gene upregulation and distinct compensatory changes in other Eph/ephrin family members, including genes previously associated with metastatic progression. Because these compensatory responses differed between models and were not recapitulated *in vitro*, they likely reflect tumor microenvironment-dependent feedback mechanisms rather than uniform cancer cell-intrinsic responses. Although the effects of EPHA4 loss differed quantitatively between the models, increasing metastatic colony size in MDA-MB-231 cells and colony number in 4T1 cells, the overall directional phenotype toward increased metastasis was consistent.

This study has several limitations. First, our mechanistic analysis focused primarily on EPHA4 function in primary tumors and early metastatic phenotypes, while its role in fully established metastatic lesions remains unclear. Second, although our data support a tumor-intrinsic role for EPHA4 in regulating epithelial gene expression, the signaling mechanisms linking EPHA4 activity to transcriptional control of cell state remain unresolved. Addressing these questions will be important for elucidating how EPHA4 integrates into broader EMP regulatory networks. While our study does not directly establish the mechanistic basis of EPHA4-dependent state regulation, these findings suggest that Eph/ephrin signaling represents an underappreciated axis influencing cancer progression through modulation of cell state transition

Overall, this work identifies EPHA4 as a regulator of mesenchymal-to-hybrid state transitions in TNBC, highlights the complications of targeting pathways that control cancer cell plasticity during metastatic progression, and points to new directions for investigating the role of EPHA4 in metastasis and therapeutic resistance.

## Materials and methods

### Cell culture

Human TNBC cell line MDA-MB-231 (MDA231), human embryonic kidney epithelial cell line HEK293T, and mouse TNBC cell line 4T1 were purchased from American Type Culture Collection (ATCC). All cell lines are routinely tested for Mycoplasma contamination using the MycoAlert kit (Lonza, LT07-218) and used within 10 passages. MDA-MB-231 and HEK293T were cultured in DMEM media (Gibco, 11965-118) supplemented with 10% fetal bovine serum (FBS, Avantor, 97068-085), 1% penicillin/streptomycin (Gibco, 15140-122). 4T1 cells were cultured in RPMI1640 media (Gibco, 11875119) supplemented with 10% FBS and 1% penicillin/streptomycin. All cells were grown in a humidified incubator with 5% CO_2_ at 37 °C.

### Generation of EPHA4 knockout cell lines

EPHA4 knockouts (KO) were generated in MDA-MB-231 and 4T1 cells by introducing frameshift indel mutations using the CRISPR-Cas9 system via lentiviral transduction. Single guide RNAs (sgRNAs) targeting either human or mouse EPHA4 were cloned into the lentiCRISPR v2 plasmid (Addgene #52961) following the protocol established by the Feng Zhang laboratory.

The following sgRNA sequences were used: human EPHA4 (5’-TGGAGTAGCCCTCAGAATAC-3’) and mouse EPHA4 (5’-TACACCGATCCGAACCTACC-3’). Both sgRNAs were verified in silico to target all annotated mRNA variants using the GRCh38 (human) and GRCm39 (mouse) reference genomes. Plasmids were amplified by overnight culture in NEB® 5-alpha E. coli (NEB, C2987H) and purified using the HiSpeed Plasmid Midi Kit (Qiagen, 12643). Lentiviral particles were produced by transfecting HEK293T cells (20-30% confluency in a T-25 flask) with a mixture containing Opti-MEM (240 µL, Gibco, 31985-070), pCMV-VSV-G (1.6 µg), pMDLg/pRRE (2.56 µg), pRSV-Rev (2.56 µg), lentiCRISPR-sgRNA vector (3.4 µg), and LT-1 transfection reagent (22.8 µL, Mirus, MIR2305). Cells were incubated for 72 hours post-transfection, after which the viral supernatant was collected and filtered using a 0.45 µm PES syringe filter. MDA-MB-231 and 4T1 parental cell lines were exposed to a mixture of 1 mL culture medium, 1 mL filtered viral supernatant, and polybrene (10 µg/mL) for 72 hours to achieve lentiviral transduction. Selection of successfully transduced cells was achieved by culturing in medium containing puromycin (2 µg/mL for MDA-MB-231; 4 µg/mL for 4T1) for at least 1 week, until non-transduced cells were eliminated. The resulting polyclonal populations were maintained in regular culture medium. EPHA4 protein knockout was confirmed by western blot analysis.

### Mouse studies

All mouse studies were conducted following protocols approved by the University of Chicago Institutional Animal Care and Use Committee (IACUC #72228). BALB/c mice (Charles River, 028) and athymic nude mice (Charles River, 490) were acquired through the Animal Resources Center at the University of Chicago. Mice were randomized before being assigned to an experimental group.

For the bulk RNA-seq study, 2 × 10^6^ MDA-MB-231 control or EPHA4 KO cells were resuspended in 100 µL 1:1 (v/v) PBS : Basement Membrane Extract (Cultrex, 3632-010-02) and orthotopically injected into the mammary fat pad of 6-week-old, female athymic nude mice. Tumor growth was monitored weekly by caliper measurement. Tumor volume was calculated by the equation: Volume = 0.4 × length × width^2^. Mice were euthanized 10 weeks post cancer cell injection, and tumor weight was recorded. Tumors were divided into 3 parts for RNA extraction (snap-frozen), protein extraction (snap-frozen), and histology (formalin-fixed, paraffin-embedded, or FFPE).

For determining 4T1 primary tumor growth and spontaneous metastasis, 2 × 10^4^ 4T1 control or EPHA4 KO cells were resuspended in 100uL PBS injected into the mammary fat pad of 6-week-old, female BALB/c mice. Tumor growth was monitored biweekly. Mice were euthanized 6 weeks after injection. Overt surface metastases in the lungs were counted after perfusion and formalin fixation. Fixed lungs were sectioned in 5 um thickness for H&E staining. The internal metastasis burden was quantified by (total number of cancer cells) / (total number of cells in the section) × 100%. The number and size of metastatic colonies were also recorded.

To assess the ability of cancer cells to colonize the lung directly, 2 × 10^6^ MDA-MB-231 cells or 8 × 10^4^ 4T1 cells were injected into the tail vein of athymic nude or BALB/c mice as indicated. Mice were monitored twice per week for body weight changes and signs of discomfort. 14 days (4T1) or 8 weeks (MDA-MB-231) post injection, the mice were euthanized. Metastasis burden in the lung was measured for surface and internal metastasis as described above.

### Bulk RNA-seq analysis of MDA-MB-231 tumors

Snap-frozen MDA-MB-231 control and EPHA4 KO primary tumors were placed in M Tubes (Miltenyi Biotec, 130-093-236), and homogenized in TRI Reagent (Zymo Research, R2050-1-200) by the gentle MACS Dissociator (Miltenyi Biotec 130-093-235, program h_RNA_02).

Genomic DNA was removed, and total RNA was extracted using the Direct-zol RNA MiniPrep Plus kit (Zymo Research, R2072) following manufacturer’s instructions. RNA concentration measurement, quality control, library preparation, and sequencing were all performed by the University of Chicago Genomics Facility following standardized protocols. Specifically, all samples were confirmed with RNA integrity number (RIN) between 8.20 - 9.50 and sequenced on the Illumina NovaSeq X instrument, using the paired end, 100bp (PE100) format and targeting 50-60M reads per sample.

Raw RNA-seq data was processed by the University of Chicago Bioinformatics Core with a standard analysis pipeline. Briefly, the raw sequencing data in FASTQ was preprocessed with FastQC for quality check and adaptor trimming. The reads were then aligned to GRCh38/GRCm39 human/mouse genome assembly using the STAR tool (Spliced Transcripts Alignment to a Reference)(26), and the mapping quality was assessed using Picard (Broad Institute). Transcript abundances were then converted into a raw count matrix using the Salmon pipeline, and the raw count matrix was imported into DESeq2 to facilitate subsequent transformations and modeling. Genes expressed in two or fewer samples were excluded. CPM normalization was adjusted for differences in library size, and then the Trimmed Mean of M-values (TMM) method from *edgeR* was applied to account for composition biases between libraries. Normalized data was first transformed using *voomWithQualityWeights*(*27*), which applies variance stabilization and accounts for sample-specific quality differences, and then fitted to a linear model using *limma*(*28*). This allows the detection of differential expression between experimental conditions by computing log fold changes (LFCs) and statistical significance values. Generally, p-value <= 0.05 and absolute log2FC >= 0.58 are used as threshold parameters to obtain differentially expressed genes. Metascape was used fo functional analysis of differentially expressed genes. Gene set enrichment analysis (GSEA) was used for determining the directional changes of biological pathways.

### Preparation of single cells from 4T1 tumors

Six weeks post mammary fat pad injection, two control and two EPHA4 KO primary 4T1 tumors in BALB/c mice of similar size were dissected and dissociated into single cells. Briefly, tumors were minced and dissociated with RPMI media containing 1mg/mL Collagenase IV (Sigma-Aldrich, C5138), 0.1mg/mL Hyaluronidase V (Sigma-Aldrich, H6254), 20U/mL DNase I Type IV (Sigma-Aldrich, D5025) with gentle shaking at 37 °C for 30 min. Cell suspensions were passed through a 70 um strainer, washed with RPMI media, and centrifuged at 300 g for 5 min. Cell pellets were resuspended with 1x Red Blood Cell Lysis Buffer (Miltenyi Biotec, #130-094-183), incubated at 37 °C for 2 min and neutralized with RPMI media. Cells were filtered through a 30um strainer and centrifuged again. Dead cells were removed using the Dead Cell Removal Kit (Miltenyi Biotec, #130-090-101) according to the manufacturer’s instructions. Cell viability > 85% was confirmed with Trypan blue staining. Cell suspensions were immediately submitted for cDNA library preparation using the Chromium GEM-X Single Cell 3’ Kit v4 (10× Genomics, 1000686) according to manufacturer guidelines. Libraries were sequenced on the Illumina sequencer at the University of Chicago Genomics Facility.

### Single cell RNA-seq analysis of 4T1 tumors

#### Preprocessing and quality control

scRNA-seq raw data was demultiplexed using the mkfastq function from the 10x Genomics Cell Ranger pipeline (version 8.0.1) to generate FASTQ files, which were then aligned to the GRCh38 (2020-A)/GRCm10(2020-A) reference genome. Low quality reads were filtered out using the default method. Quality control was performed based on the following criteria: (1) Cells with fewer than 200 detected genes or genes expressed in fewer than 3 cells were removed (2) Cells with >20% mitochondrial gene expression were filtered out, as this is often indicative of compromised cell viability or apoptosis (3) Doublets were identified and removed using DoubletFinder (version 2.0.3). The gene expression matrix was then log-normalized using the NormalizeData function in Seurat (version 4.3.0, (29)) to account for differences in sequencing depth across cells. Highly variable genes (HVGs), which are genes that exhibit significant variation across cells and are likely to be biologically relevant, were identified using the FindVariableFeatures function. The top 2,000 HVGs were selected for downstream analysis to reduce computational complexity while retaining biologically meaningful signals. Linear dimensionality reduction via integrated canonical correlation analysis (CCA) was used for dimensionality reduction to compute a shared clustering representation across experimental conditions to enable comparison.

#### Cell clustering and cell type annotation

We constructed a k-nearest neighbors (kNN) graph based on Euclidean distances in the integrated CCA dimensionality-reduced space using the first 50 dimensions. To identify cell populations, Louvain clustering was applied on the kNN graph and a clustering resolution of 0.55, generating 27 clusters. The optimal resolution was selected by assessing clustering stability across different pairwise combinations of resolutions using the Adjusted Rand Index (ARI): the lowest resolution was selected at which categorical assignments were stable. The clusters were visualized using UMAP (Uniform Manifold Approximation and Projection) (30). To annotate tumor cells, we applied Single CEll Variational ANeuploidy analysis (SCEVAN) which discriminates between tumor and non-tumor cells by clustering cells based on copy number variation (CNV)(31). Tumor cells were defined as cells designated as tumor by SCEVAN and had zero Ptprc expression. Cells within the main tumor cell clusters (clusters 0, 1, 4, 6, 10, 14 and 21) that were not designated as tumor cells by SCEVAN, as well as cells annotated as tumor cells by SCEVAN that were presented in non-tumor clusters, were excluded from subsequent analysis due to unclear identities. The remaining cells were annotated manually by examining cluster-wise expression of cell type marker genes from a previous scRNA-seq study (32). Clusters were assigned a cell type if they expressed the marker genes for that cell type more than other clusters. After all annotation steps, clusters 7, 9, 13 and 25 remained unannotated as they did not highly express any markers for expected TME cell types and were excluded from subsequent analysis.

#### Sub-clustering and differential gene expression analysis of 4T1 cancer cells

Using the tumor cells as defined above, we applied linear dimensionality reduction via integrated CCA and constructed a kNN graph using the first 9 dimensions. We applied Louvain clustering on the kNN graph using a clustering resolution of 0.25, optimized using ARI, generating 5 clusters representing different tumor cell states.

To determine the molecular signature for each cell state, we performed differential gene expression analysis using the FindAllMarkers function in Seurat. The top positive and negative cluster markers (adj.p-value < 0.05, log2FC > 1, pct.1 > 0.25 for positive markers or adj.p-value < 0.05, log2FC < -1, pct.2 > 0.25) were analyzed with Metascape (33) using the default setting to determine the enriched functions. To identify EPHA4 regulated functions within each cell state and across all cell states, we used the FindMarkers function in Seurat to identify genes differentially regulated in EPHA4 KO relative to control tumor cells. The top up-regulated (adj. p-value < 0.05, log2FC > 0.5, pct.1 > 0.1) and down-regulated genes (adj. p-value < 0.05, log2FC < -0.5, pct.2 > 0.1) were analyzed with Metascape using the default setting to determine the enriched functions.

#### Gene signature score calculation for single cells

Gene signature scores were calculated using the AddModuleScore function in Seurat. The epithelial signature included *Ovol2, Grhl2, Esrp1, Ovol1, Ocln, Cldn7, Cldn4, Cldn3, Epcam, Esrp2,* and *Cdh1*, and the mesenchymal signature included *Zeb1, Zeb2, Twist1, Twist2, Snai2, Vim, Fn1, Acta2,* and *Cdh2*. Hallmark mouse gene sets for cell state annotation and quantification of biological programs were obtained from the GSEA MSigDB Mouse Collections.

#### Cell-cell communication inference

Cell-cell communication networks were inferred using the CellChat R package (14) to investigate interactions between cell types and assess the impact of EPHA4 KO on EPHA and EPHB signaling pathways. Interaction analyses were performed using 11 defined tumor microenvironment (TME) cell types, as well as 5 clusters identified within the cancer cell population. CellChat quantifies ligand-receptor mediated communication between pairs of cell groups based on the average expression levels of ligands in one group and corresponding receptors (including relevant cofactors) in another group. Statistical inference was then applied to identify biologically significant interactions.

### Boyden chamber invasion assay

Inserts (Corning, 353097, 8.0 um) were plated in 24-well plates (Corning, 353504) and coated with 200uL 0.25mg/mL basement membrane extract (Cultrex, 3445-010-01) diluted in DMEM (MDA-MB-231) or RPMI (4T1) media supplemented with 1% Pen/Strap and incubated for 1 hour at 37℃ to solidify. Control and EPHA4 KO cells were digested, neutralized with media with 10% FBS, and resuspended in serum-free media. 1 × 10^5^ MDA-MB-231 or 4T1 cells in 300uL serum-free media were added to the upper chamber of each insert. 500uL DMEM or RPMI media with 10% FBS and 1% Pen/Strep was added to the lower chamber as chemoattractant. Serum-free media was added to additional wells with inserts as negative controls. Cells were allowed to invade at 37℃ for 16 - 24 hours. To quantify the invaded MDA-MB-231 cells, cells were stained with 4ug/mL Calcein AM (Corning, 354216) in HBSS buffer for 1 hour. Uninvaded cells in the upper chamber were removed by swabbing with cotton tips. The invaded cells were dissociated from the bottom side of the insert by incubating in 500uL cell dissociation buffer (Gibco, 13150016) with gentle agitation at 140 rpm for 1 hour. The Calcein AM signal was measured using a Perkin Elmer Victor X3 plate reader as a read-out of invaded cells.

Because 4T1 cells are sensitive to serum deprivation, to quantify invaded 4T1 cells, the inserts were directly fixed in 2% paraformaldehyde/PBS at room temperature for 1 hour after removing uninvaded cells from the top chamber. The invaded cells were stained with Hoechst staining at 1:1000 dilution for 15 minutes at room temperature. Invaded cells were counted in 20 randomly selected fields by fluorescence imaging, and the average number of cells in each field was calculated and compared across conditions.

### RNA isolation, reverse transcription and qRT-PCR

Cells were directly lysed in RLT buffer with 1% v/v 2-Mercaptoethanol. RNA was isolated using RNeasy Mini Kit (Qiagen, 74106). 4 µg of total RNA from each sample was converted to cDNA using High-Capacity cDNA Reverse Transcription Kit (Applied Biosystems, 4368813). cDNA was diluted 10 - 20 times in ddH2O. Standards were prepared by pooling cDNA from all samples followed by serial dilution of 1:10, 1:100, and 1:1000 fold. Sample and standard cDNA were mixed with SYBR Green Master Mix (Applied Biosystems, A25742) and analyzed on LightCycler® 96 Instrument (Roche). The expression level of genes was normalized to RPL13A. Primer pairs used for this study:

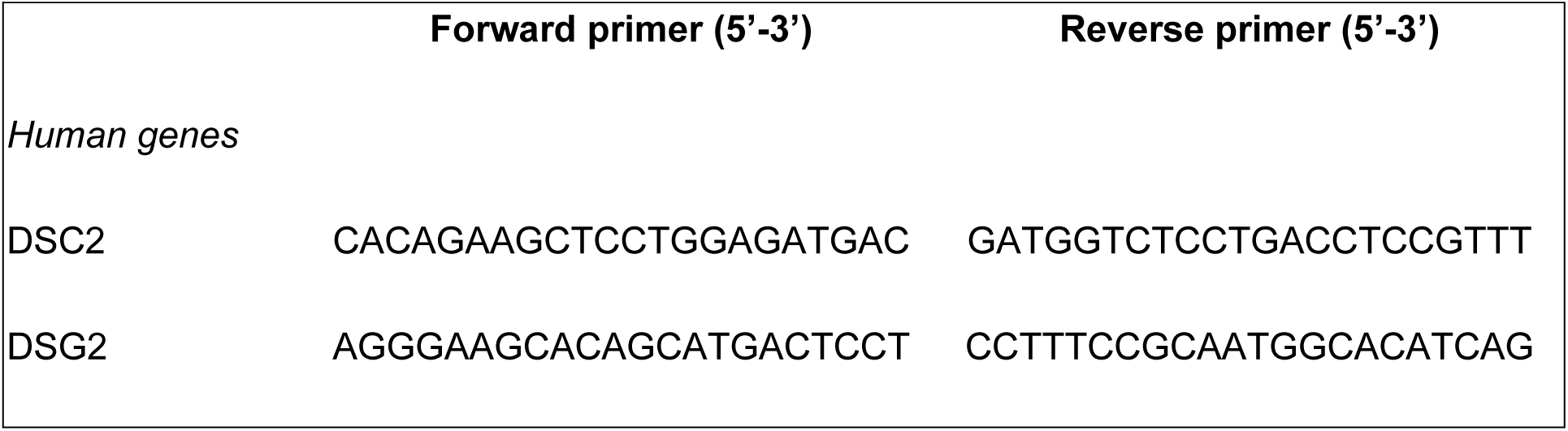
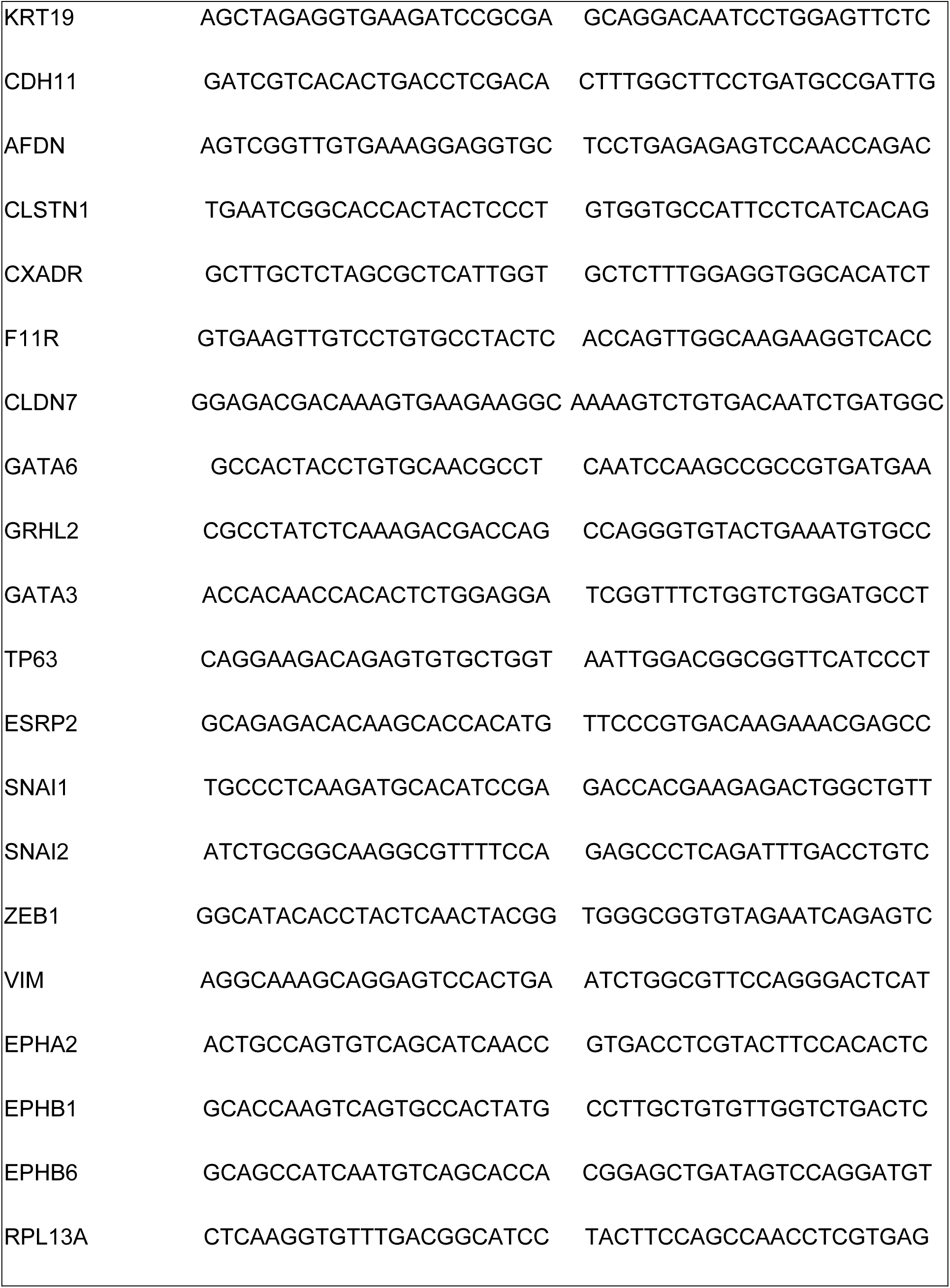
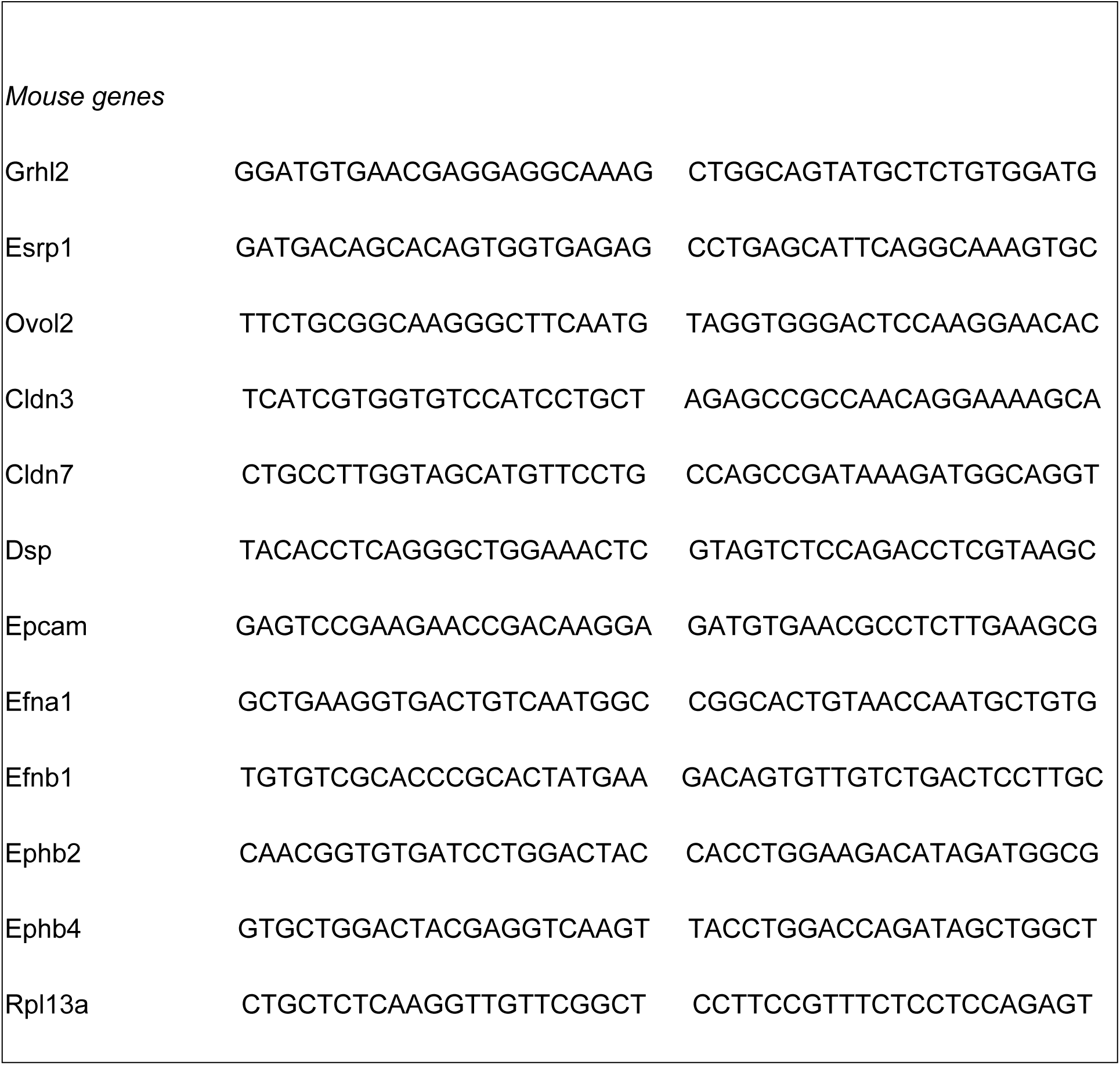

### 3D cell culture

For 3D culture, 1 × 10^4^ 4T1 cells or 2.5 × 10^4^ MDA-MB-231 cells were resuspended in 30uL Matrigel (Growth factor reduced, Corning, 356230, thawed at 4℃ overnight) and plated into 24-well, flat bottom, low attachment plate (Corning, 3473) as a dome. The Matrigel dome was solidified at 37℃ for 30 minutes before adding complete media into the well. After 72 hours of culture, culture media was removed. The Matrigel dome was broken up by gentle pipetting in 500uL Cell Recovery Solution (Corning, CLS354253). The Matrigel-cell suspension was then transferred to an Eppendorf tube and incubated on ice with gentle agitation for 30 minutes. Cells were pelleted by centrifuging at 500g for 5 minutes at 4℃ and lysed in 350uL RLT buffer for RNA isolation.

### Histology staining of FFPE slides

Tumors and lungs were fixed in Bouin solution (Sigma Aldrich, HT10132-1L) and stored in 70% ethanol. Paraffin embedding, cross-sectioning, and histology staining was performed by the University of Chicago Human Tissue Resource Center following standard protocols. One slide of each sample was stained with the hematoxylin and eosin stain. Stained slides were scanned by Olympus VS200 Slideview scanners at 40x magnitude at the Integrated Light Microscopy Core and then analyzed with QuPath software.

### Western blotting

Cells cultured in 6-well dishes were directly lysed in 100uL Laemmli Sample Buffer (Bio-Rad, 1610747), boiled at 95℃ for 5 min and sonicated for 10 seconds at 15% amplitude. Protein lysates were loaded on 4-15% Criterion™ TGX™ Precast Midi Protein Gels (Bio-Rad, 5671084) and separated by electrophoresis at 120 V for 1 hr. Proteins were transferred to PVDF membrane (EMD Millipore, IPFL00010) at 4℃, 30 mA for 1.5 hours. The membranes were blocked in 5% milk or 3% BSA at room temperature for 1 hour. Primary antibodies were added, and the membranes were incubated at 4℃ overnight with gentle agitation. On the second day, the membranes were washed 3 times in TBST buffer, then incubated with secondary antibodies for 1 hour at room temperature. The membranes were washed 3 times in TBST buffer, incubated with Pierce™ ECL Western Blotting Substrate (Thermo Fisher, 32209) for 2 minutes, and imaged with LI-COR Image Studio Software.

The following antibodies were used in this study: Anti-αTubulin mouse monoclonal antibody (Invitrogen #MA1-19401), 1:1000 in 3% BSA; Anti-EPHA4 mouse monoclonal antibody (Invitrogen #37-1600), 1:500 in 5% milk; Anti-GAPDH mouse monoclonal antibody (Santa Cruz Biotechnology #sc-32233), 1:1000 in 5% milk; Goat anti-Mouse IgG-HRP secondary antibody (Invitrogen, #31430, 1:3000 dilution in TBST buffer)

### Analysis of breast cancer public datasets

We used the following public datasets to analyze gene expression in patients and cell lines: (1) The Cancer Genome Atlas (TCGA) Firehose Legacy breast cancer dataset on cBioportal for gene expression in primary breast tumors, (2) ISPY2 breast cancer dataset for gene expression in high risk, early stage patients (GSE194040) (3) GSE110590 for gene expression in matched breast primary tumor and metastasis sites, (4) DepMap for gene expression quantification in cancer cell lines. Detailed analytical procedures were described in the relevant figures.

## Acknowledgements

We sincerely thank Margarite M. Kohs, Joana Pinhero, Kent Schecter, Emily Shi for their valuable discussions and insightful contributions. We acknowledge the University of Chicago Human Tissue Resource Center (RRID: SCR_019199) for histology staining, the University of Chicago Genomics Facility (RRID: SCR_019196), especially P. Faber for assistance with RNA-seq, and the University of Chicago cell screening center. This work was supported by U.S. Department of Defense (W81XWH2110120) (M.R.R.). We acknowledge the Susan G. Komen TREND Grant (TREND21675016) (W.L.), the Goldblatt Endowment Fund (W.L.), the Fitch Scholarship Fund (W.L.), philanthropic support for the Cancer Health Equity Training Program (W.L., L.C.N.), the Biological Sciences Collegiate Division Research Endowments at the University of Chicago (H.W.L., T.L.), UChicago Quad Undergraduate Scholars Fund (T.L.), Susan G. Komen Career Transition Award (L.C.N.), and the University of Chicago Comprehensive Cancer Center (UCCCC) Sigal Fellowship (L.C.N.).

## Author Contributions

W.L. and M.R.R. conceptualized the study. W.L., H.W.L., M.H.B., L.C.N., and M.R.R. contributed to methodology development. W.L., H.W.L., M.H.B., L.C.N., A.K., T.J.L., G.P.Y., and E.S. performed the investigation. H.W.L., G.P.Y., and W.L. conducted bioinformatics analysis, curated the data and generated visualizations. W.L. and M.R.R. managed project administration.

M.R.R. provided resources, supervision, and funding acquisition. W.L. and M.R.R. wrote the original draft of the manuscript, and H.W.L. contributed to the Methods section. All authors reviewed the manuscript and provided feedback. M.R.R. and W.L. revised and edited the manuscript. For correspondence: email M.R.R at.

## Data availability

All data generated or analyzed during this study are included in this published article and its Supplementary Information files, unless otherwise stated. Bulk RNA-sequencing and single-cell RNA-sequencing datasets generated in this study has been deposited in the NCBI Gene Expression Omnibus (GEO) (MDA-MB-231 bulk RNA-seq: GSE333898; 4T1 scRNA-seq: GSE333901), and will be made public upon acceptance of the manuscript. Publicly available datasets analyzed in this study include The Cancer Genome Atlas (TCGA) Breast Invasive Carcinoma cohort, accessed via cBioPortal, and breast cancer cell line expression data from the DepMap Portal.

