## Supplementary figures and images for "EPHA4 Regulates the Transition between Mesenchymal and Hybrid States in TNBC"

### Supplemental figures

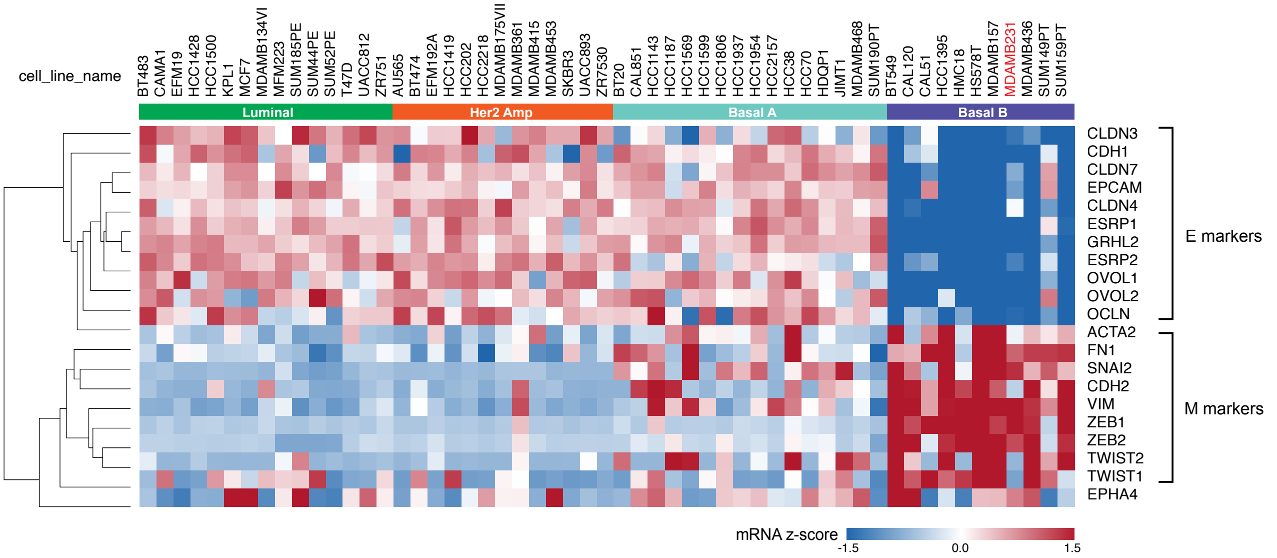

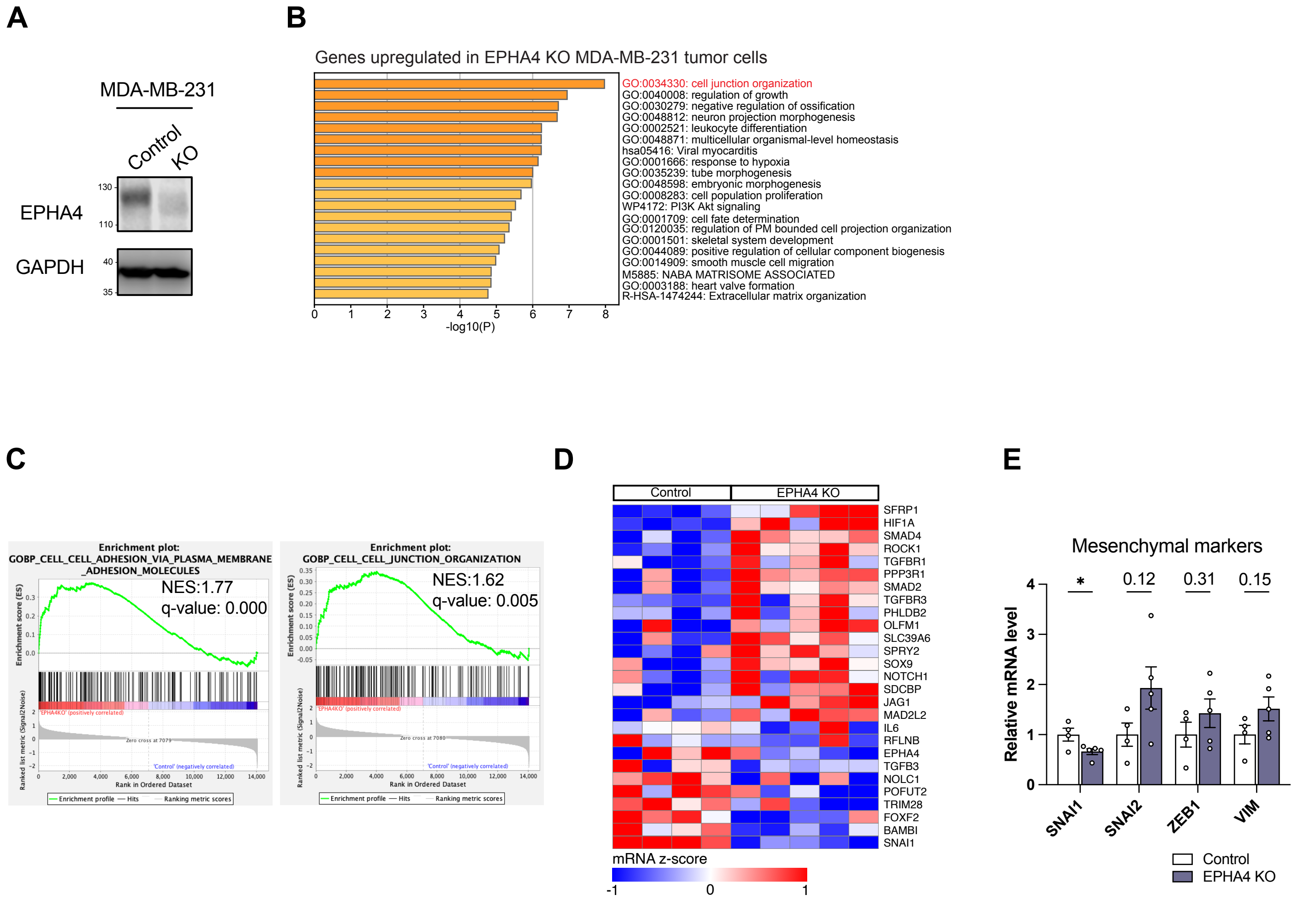

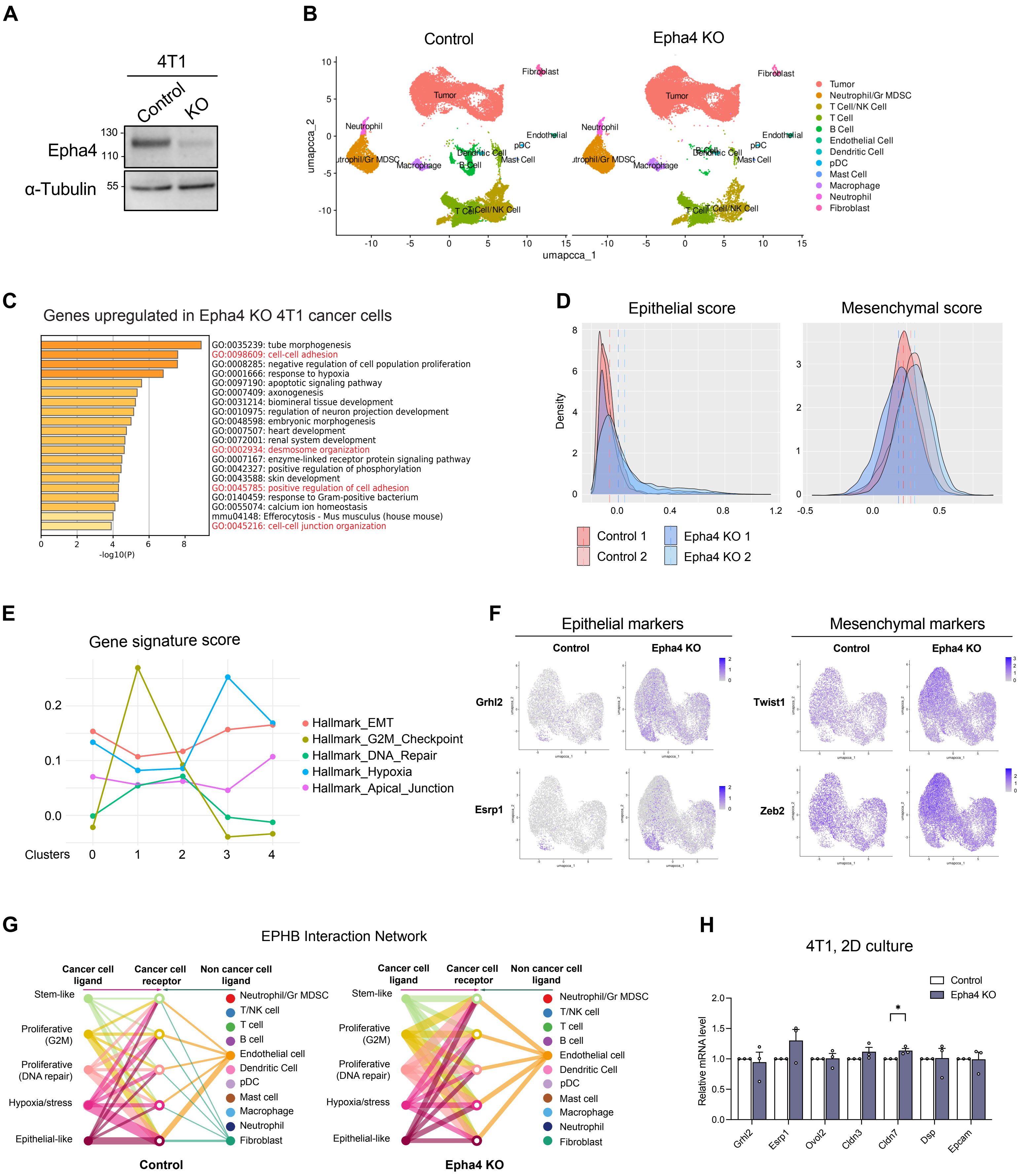

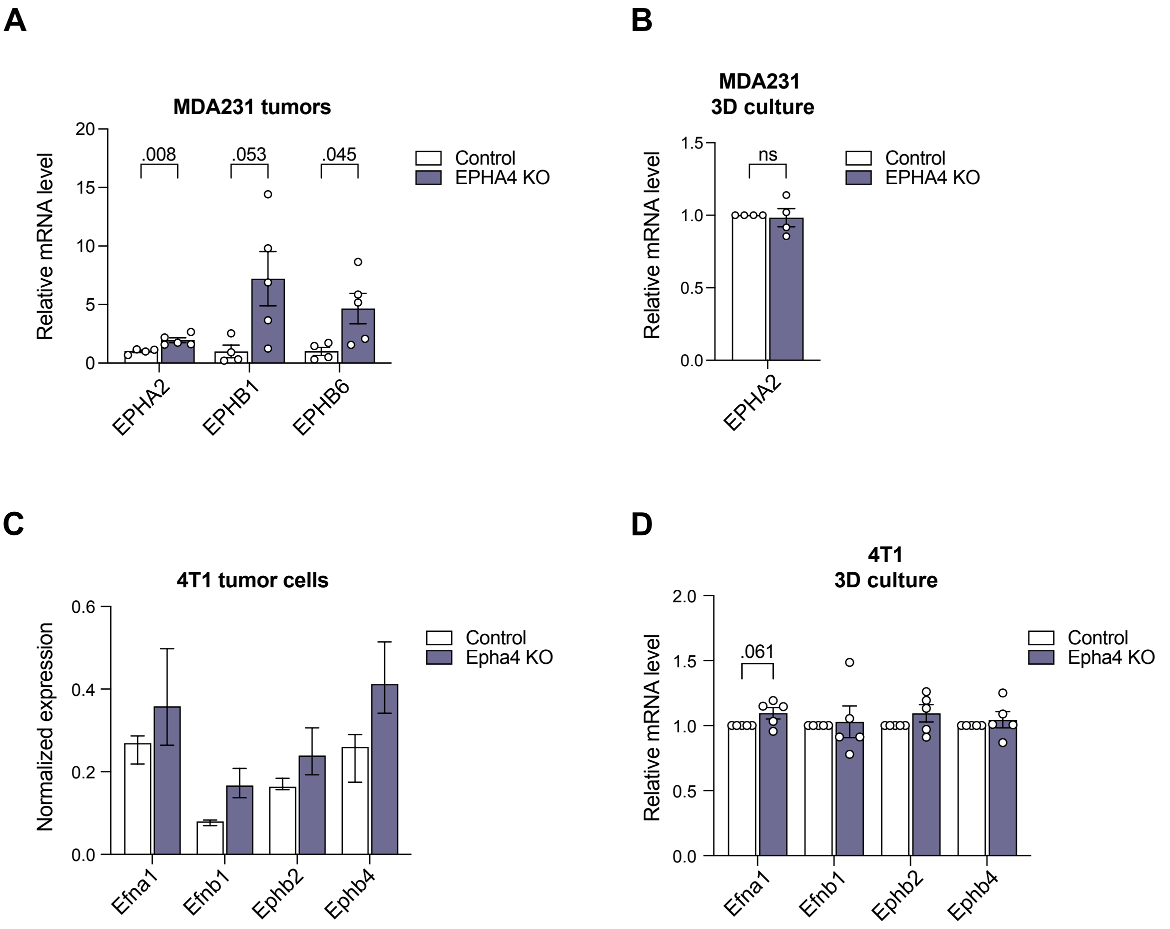
